# Dynamic regions allosterically connect the USP14 active site with the proteasome interaction surface

**DOI:** 10.1101/2024.06.28.601165

**Authors:** Johannes Salomonsson, Linda Sjöstrand, Arvid Eskilson, Dean Derbyshire, Pádraig D’Arcy, Maria Sunnerhagen, Alexandra Ahlner

## Abstract

<u>U</u>biquitin-<u>s</u>pecific <u>p</u>rotease 14 (USP14), is a member of the USP family responsible for the catalytic removal of ubiquitin (Ub) from proteins directed to the proteasome, implicated in the pathogenesis of neurodegeneration and cancer. Crystallography and cryo-EM analysis have identified loop regions crucial for the deubiquitinase activity of USP14, specifically those involved in Ub and proteasome binding. However, the structural changes in USP14 upon ligand binding to these regions are minimal, indicating significant yet uncharacterized dynamic contributions to its function. In this study, through structural and dynamical NMR experiments and functional evaluation, we demonstrate that small mutations designed to impact Ub binding and catalytic activity without disturbing the USP structure display both local and long-range effects. The affected residues connect the active site and the Ub binding region with the proteasome interaction surface through a network of loops, which show varied dynamics on the ps-ms time scale. Collectively, our findings experimentally reveal different aspects of dynamic connections within USP14, suggesting the presence of allosteric networks that link enzyme activity with regulatory function. The novel concept that USP14 allosteric networks are pre-existing, coupled, and activated by regulatory interactions with the USP fold, could be crucial to future targeted drug design.

## Introduction

The ubiquitin-proteasome system (UPS) plays a crucial role in maintaining protein homeostasis by coordinating the degradation of damaged, misfolded, or short-lived proteins. At its core is the proteasome, a large multisubunit proteolytic complex functioning as the cell’s molecular shredder, and ubiquitin (Ub)—a highly conserved 8 kDa protein serving as a specific degradation tag. Ubiquitination, involving E1, E2, and E3 ligases, sequentially attaches Ub to target proteins, forming mono- or polyubiquitin chains. Conversely deubiquitinases (DUBs) counter this by removing Ub from target substrates [1,2]. Humans have approx. 100 DUBs spanning seven families, categorized by sequence and structure homology. The largest and most diverse is the ubiquitin-specific protease (USP) family, with over 50 members, all sharing a conserved USP domain responsible for Ub binding and catalytic activity [3,4].

USP14, one of three proteasome-associated DUBs, regulates protein degradation by removing ubiquitin from substrates and modulating proteasome activity through allosteric mechanisms [5–7]. Although it exhibits limited DUB activity in isolation, its catalytic efficiency is enhanced upon association with the proteasome [8,9]. USP14 comprises two distinct domains, connected by a flexible linker: a ubiquitin-like (Ubl) domain and a USP domain responsible for its catalytic activity. Upon proteasome binding, the Ubl domain of USP14 (USP14_Ubl_) anchors to the proteasome PSMD2 subunit (also known as Rpn1 in yeast), while its catalytic USP domain interacts with the PSMC1 (Rpt2) and PSMC2 (Rpt1) subunits [9,10]. Recent cryo-EM studies reveal variations in USP14’s interaction with the proteasome suggesting a model for USP14’s allosteric regulation of proteasome motion and function [11].

The USP14 USP domain (USP14_USP_) features a conserved open hand-like fold, with the protease active site situated in a groove between the “thumb” and “palm”. Upon binding, the Ub moiety is positioned in the “palm”, supported by the Ub-binding loops within the USP fold [3,12]. USP14_USP_ has been characterized structurally by crystallography in its free state (PDBID: 2AYN), when bound to Ub (PDBID: 2AYO) [12], when inhibited by small molecules (PDBID: 6IIK, -N, -L, -M) [13], and as a catalytically inactive mutant (PDBID: 6LVS; C114S) [14]. The core structure of the USP14_USP_ [12] remains well resolved and highly consistent across USP14 structures. This includes the catalytic triad (C114, H435, D451), where USP14 appears to be properly aligned for catalysis both in the presence and absence of Ub or proteasome [11–13]. Interestingly, this contrasts to inactive “apo” structures of closely related USP7 and USP15, where the catalytic triad is misaligned [15,16]. The prevailing hypothesis for USP14 autoinhibition is based on its 3.5 Å apo crystal structure, where surface blocking loops named BL1 and BL2 partially occupy the active site groove, presumably blocking access to the catalytic site. (PDB-ID: 2AYO; [12]). The enzymatic activity of free USP14 is significantly enhanced by phosphorylation of S432 in BL2, which is replicated by the phosphomimic mutation S432E suggesting a charge-induced BL2 loop opening to relieve autoinhibition [8], however, no structures have confirmed this mechanism.

Recent cryo-EM studies have provided novel insights into USP14 as a multistate allosteric regulator on binding to the proteasome. In these structures, USP14 interacts with the proteasome subunits PSMC2-AAA and PSMC1-OB domains through a series of surface loops, many of which are unresolved in crystal structures [11,17,18]. PSMC2-AAA predominantly interacts with USP14 residues 371-391, a region disordered in crystal structures [12,13], and the USP14 PKL loop [11,18]. PSMC1-OB unit also forms several direct contacts with BL1, which in turn binds Ub, with additional PSMC1-OB touchpoints in BL2 (S430), and BL3 (W472) [11]. Although BL1 is distant from the active site, proteasome binding was proposed to position USP14 BL1 and BL2 loops for Ub binding [11] (**Figure 1C**). However, no direct connection between the proteasome binding site and the active site of USP14 was identified, suggesting allosteric activation mechanisms are at play.

**Figure 1.**
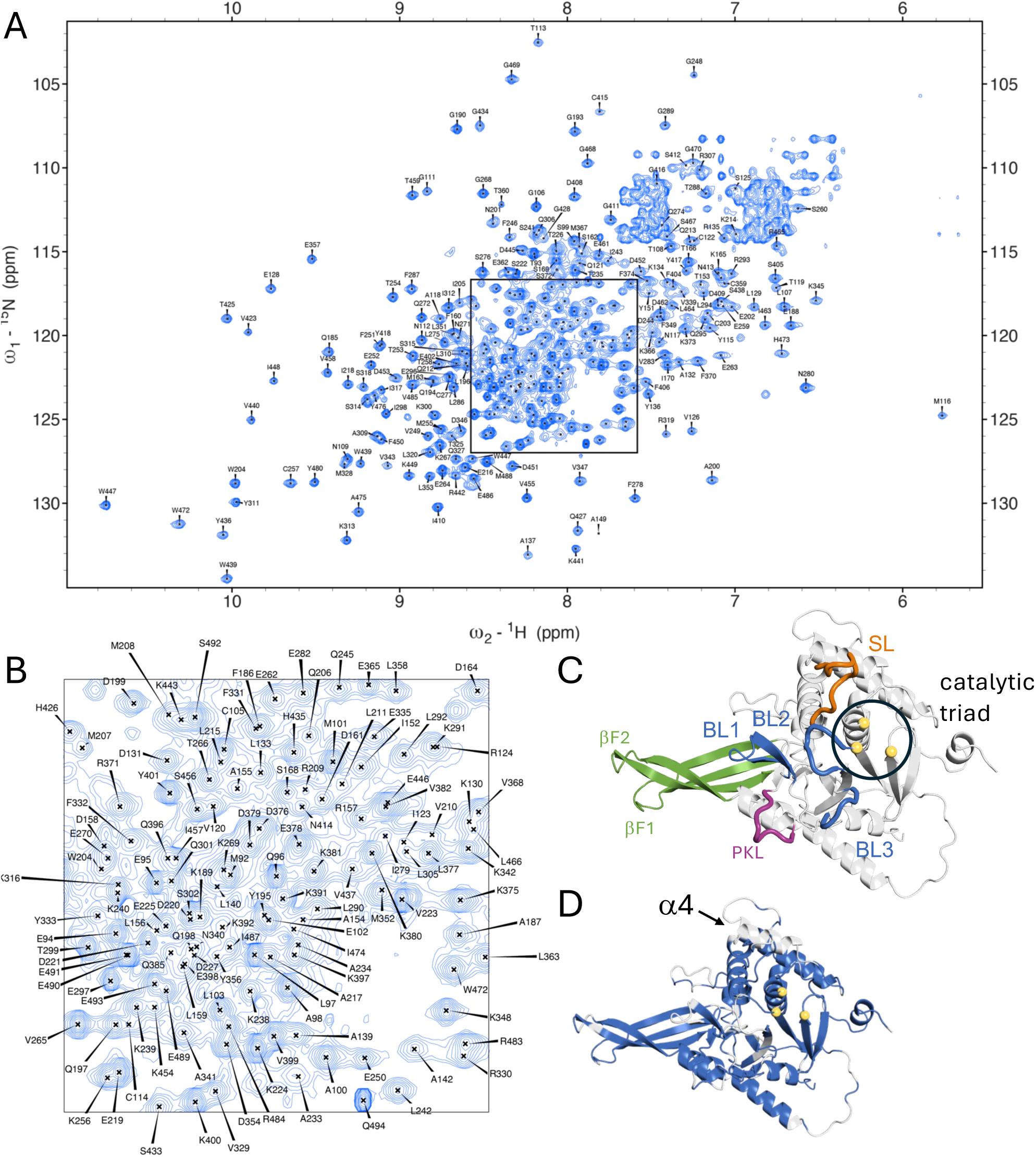
USP14 assignment, completeness and loop arrangement. (A) ^1^H-^15^N TROSY-HSQC of ^2^H-^13^C-^15^N-labeled USP14_91-494_ with assignments. (B) Magnification of the overlapped region in (A) (C) Alphafold2 model with functionally relevant loops highlighted. Switching loop (SL residues 188-199) in orange, Proximal Knuckle Loop (PKL residues 278-285) in purple, β-fingers in green (βF1 residues 249-272 and βF2 residues 295-319), and blocking loops (BL1-3) in blue (BL1 residues 330-342, BL2 residues 428-434 BL3 residues 468-473) (D) Residues with assigned chemical shifts for backbone amides are colored blue on the Alphafold2 model. Unassigned residues are shown in grey. Most of unassigned residues are in loop regions or in the core. One distinct unassigned element is helix α4 located on top of the catalytic triad residues, C114, H435 and D451 (yellow)

To fully understand proteasomal activation, there is a critical need to understand the dynamic properties of the USP14 network of functional loops in solution (BL1, BL2, BL3, PKL, SL) [19]. High-resolution structural methods such as crystallography and cryo-EM are limited in characterizing dynamic properties and provide little or no information for highly flexible regions, requiring alternative approaches[20]. By NMR, ps-ns and μs-ms dynamics can be analyzed by relaxation experiments in solution at both backbone amide and methyl groups, which has turned the technique into an excellent tool to analyze the presence of dynamically modulated function [21–26]. Added to this, the NMR chemical shift is sensitive to minute structural and/or dynamic changes[27]. In-depth experimental analysis of smaller proteins has shown the pronounced role of dynamics in allosteric networks [26,28–32]. However, due to the somewhat challenging size of the USP domain (>40 kDa), NMR studies have, to date, only been pursued for USP7, focusing on ligand and inhibitor binding [33–37], leaving the loop dynamics understudied.

To investigate the structure and dynamics of USP14_USP_ in solution we have assigned the NMR resonances of the USP14_USP_ backbone and performed relaxation analysis of its fast dynamical properties. We show that in the free state, the dynamics of USP14 is observed at a wide range of time scales, from ps to ms. Importantly, loops BL1 and BL2 near the active site are predominantly dynamic in solution in the free state. To probe whether intrinsic allosteric networks are already present in the free state of USP14, we compared NMR spectra of wild-type (WT) USP14_USP_ with those of single point mutants inhibiting (C114A) or activating (S432E) USP14 catalytic function. Additionally, we mutated residues in the USP14_USP_ Ub interaction surface. Through chemical shift perturbation (CSP) analysis we observed that functional mutations collectively impact a broader region of USP14. This region is notably conserved in USP14 and partially overlaps with the proteasome binding interface. Our work provides an extended view of different aspects of dynamics in USP14 and provides experimental evidence to support the presence of allosteric regulatory networks that connect the active site and Ub binding residues with the proteasome binding interface, thereby increasing the potential to target allosteric sites for USP14 inhibition.

## Results

### First NMR analysis of USP14_USP_ resolves dynamic loops and transient secondary structures

To study USP14_USP_ in solution, we optimized NMR conditions of ^2^H,^13^C,^15^N labeled USP14_91-494_ using TROSY-based triple resonance experiments at 30°C. The intrinsic dynamics of USP14_USP_ allowed for exchange of most of the amide protons without a denaturation/renaturation protocol, resulting in the backbone assignment of 81% of non-proline amide groups (314 of 387), 87% of C_α_, 84% of C_β_, and 85% of C’ nuclei **(Figure 1)**. Due to incomplete spin systems or weak peak intensities in the 3D experiments, 55 peaks observed in the TROSY-HSQC could not be confidently assigned.

To evaluate the structural completeness of the assignment, we mapped assigned residues onto the USP14 model from the AlphaFold Protein Structure Database (P54578) [38]. This model contains all USP14_USP_ residues, including loops (BL1-3, SL and PKL) indicated to be of importance for function [11,12,18] **(Figure 1C)** and previously unresolved regions in crystal[12] or cryo-EM structures [11]. We recently validated this USP14_USP_ model as the best fit to small-angle X-ray scattering (SAXS) data in solution [39]. Most of the unassigned residues are in surface loops or are deeply buried in the β-sheet core **(Figure 1D)**. Assignments in surface loops can be challenging due to intermediate exchange and severe spectral overlap, whereas lack of assignments for some residues located in the buried β-sheets is likely due to limited ^2^H/^1^H-exchange. Interestingly, no resonances could be assigned to residues 173-183, which according to crystal structures and the AlphaFold model form a surface-exposed α-helix (α4) located on top of the helix harboring the catalytic cysteine (α1, C114) [12] **(Figure 1D)**. Since such a highly exposed helix should not have experienced any limitations in ^2^H/^1^H exchange, it is reasonable to assume that amide resonances in α4 experience chemical exchange broadening due to millisecond dynamics.

We then proceeded to assess USP14_USP_ secondary structure in solution using chemical shift secondary structure population inference (CheSPI) [40], built on the NMR resonance assignment of H_N_, N, C’, C_α_ and C_β_ (**Figure 2A, Suppl Figure 1**). Overall, the solution secondary structure compares well to that presented in the crystal structures of the USP14_USP_ in the absence and presence of Ub (PDB-ID 2AYN, 2AYO [12]) (**Figure 2A**). An even better match was observed with the AlphaFold model (**Figure 2A**), in agreement with its excellent fit to SAXS solution data [39].

**Figure 2.**
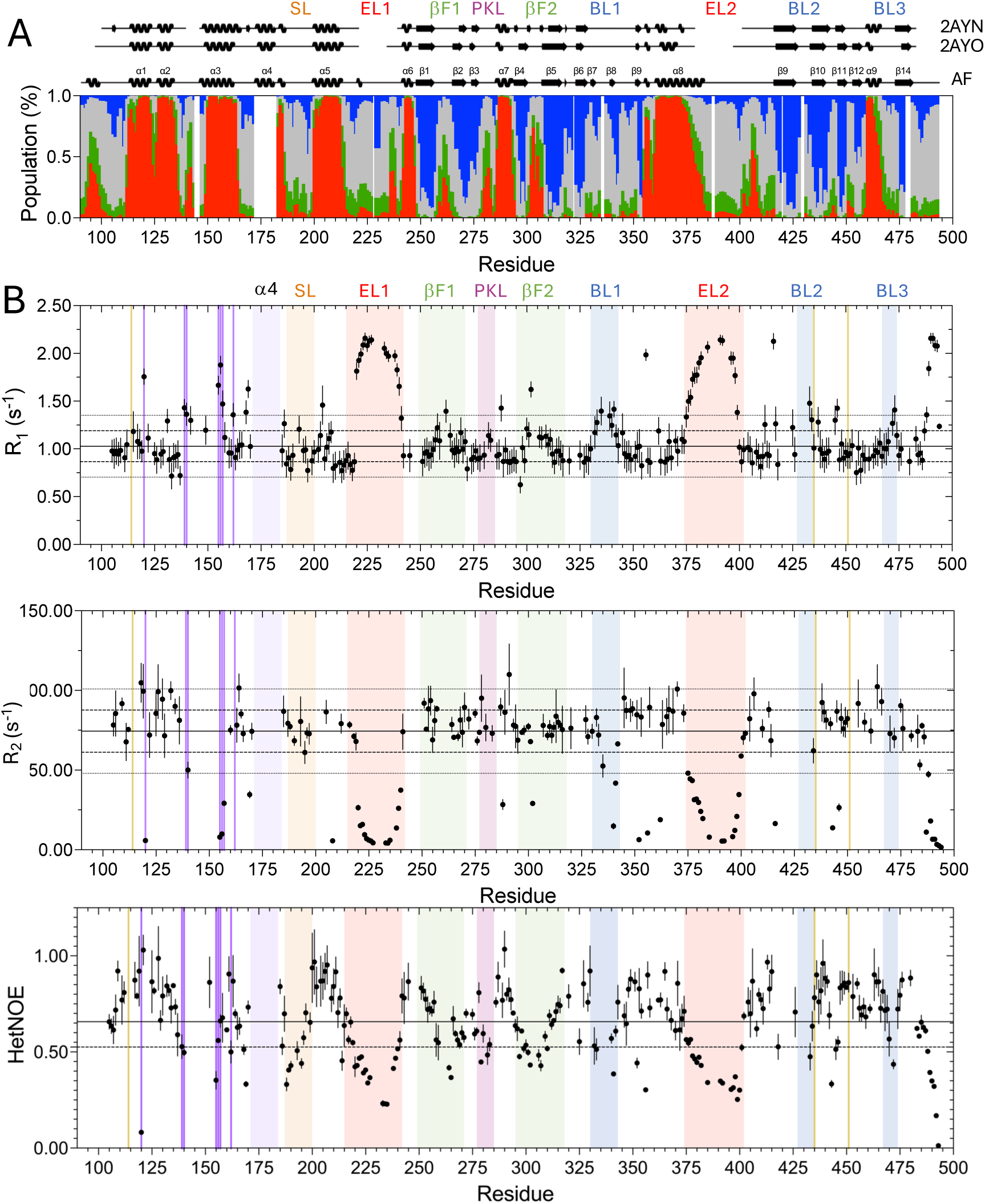
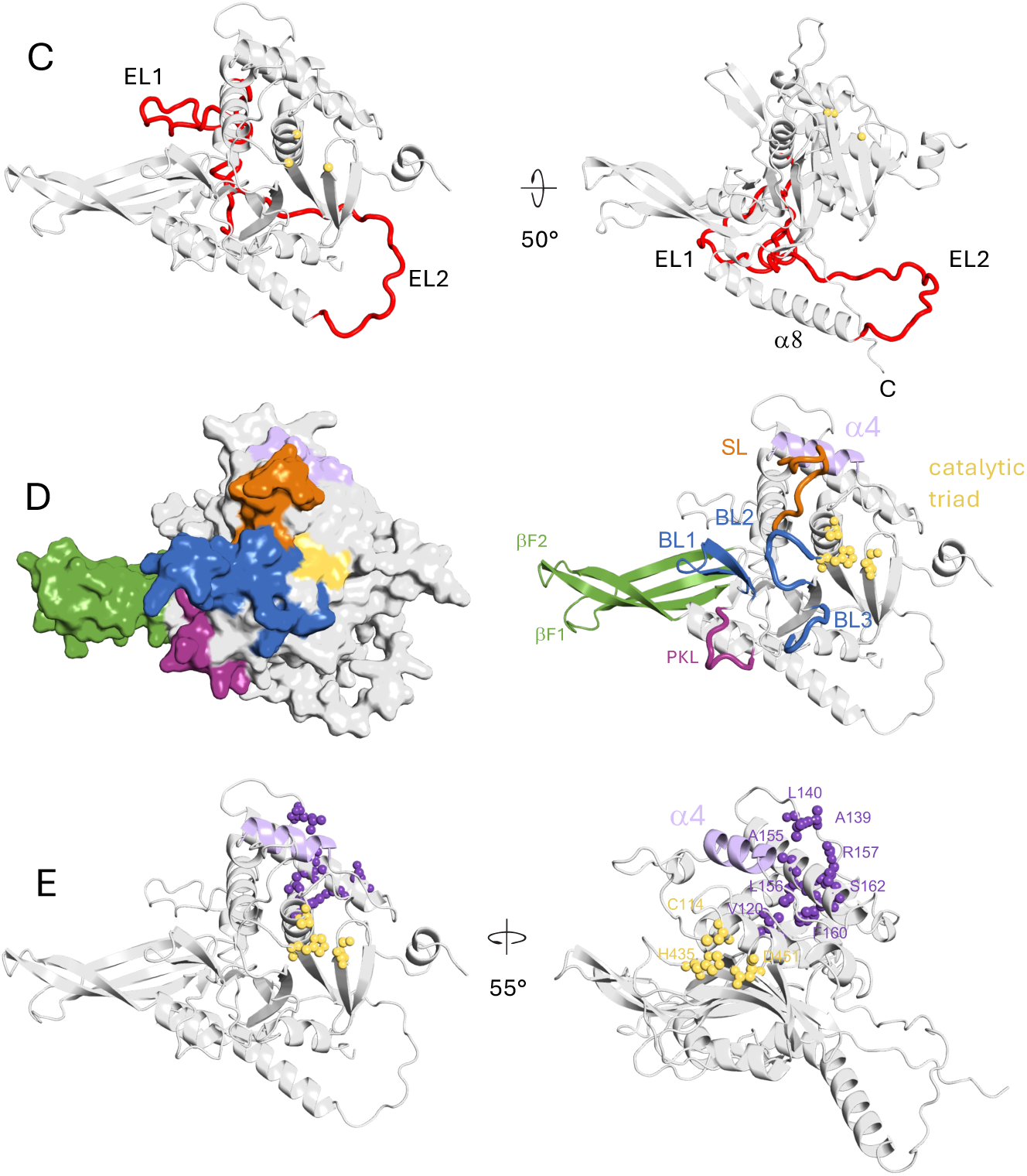
USP14 secondary structure populations and backbone NMR relaxation. Color coding throughout the figure: Helix α4 light lilac, Switching loop (SL residues 188-199) in orange, Proximal Knuckle Loop (PKL residues 278-285) in purple, β-fingers in green (βF1 residues 249-272 and βF2 residues 295-319), and blocking loops (BL1-3) in blue (BL1 residues 330-342, BL2 residues 428-434 BL3 residues 468-473), Extended loops (EL1-2) (EL1 residue 214-242 and EL2 384-416) in red, dynamic cluster in dark lilac, catalytic triad C114, H435, D451 in yellow. (A) Secondary structure calculation of USP14_USP_ from H_N_, N, C’, C_α_ and C_β_ chemical shifts derived by CheSPI [40]as a function of sequence. The populations of different structure elements are colored as extended (blue), helix (red), green (turn) and coil (grey). Secondary structure elements assigned to the crystal apo and Ub bound structures PDBID: 2AYO and PDBID: 2AYN as well as the AlphaFold2 model are indicated above the population graph, as extracted from PDBsum[68]. (B) NMR relaxation data (R_1_, R_2_ and hetNOE at 900MHz) for USP14_99-494_, revealing dynamic properties in the ps-ns range, as a function of sequence. For R_1_, R_2_ and hetNOE, the trimmed average value is indicated with a straight line. The values for one (dashed) or two (dotted) standard deviations above or below the trimmed average are indicated as lines. (C) AlphaFold model of USP14_USP_ highlighting the most flexible part of USP14_USP_ EL1 and EL2. (D) AlphaFold model of USP14_USP_ with continuous community of loops a shown as surface and cartoon view. (E) AlphaFold model of USP14_USP_ showing relaxation outliers forming an intersegmental connection between helix α4 and helix α1 containing the active C114.

By NMR, we were able to significantly contribute to improvement to existing models of two segments of USP14 that lack electron density in crystal structures, [12] **(Figure 2)** Based on this analysis, we here designate the extended loop regions EL1 (residues 214-242) and EL2 (residues 384-416). The NMR data indicate that both EL1 and EL2 lack stable secondary structure but hold transient helical turns in residues 236-239 (EL1) and 405-407 (EL2). Residues 360-383 preceding EL2 form a single, continuous helix (α8), which is stably helical until residue 375, where the helical propensity starts to gradually decline until the EL2 loop is reached. Our NMR analysis further allowed us to experimentally assess a feature of the AlphaFold USP14 structure not present in the crystal structures, where EL2 ‘extends’ beyond helix α8 wrapping around the USP14 C-terminus without forming a knot [41] **(Figure 3)**. By ^15^N-edited NOESY TROSY-HSQC experiments, we identified H_N_-H_N_ NOE cross peaks between residues 400-402 on EL2 and 485-487 in the C-terminus and could thereby verify the presence of this loop arrangement in solution **(Figure 3B)**. Together with α8, EL2 has been demonstrated to be critical for USP14 allosteric regulation of the proteasome through interactions with the AAA domain of PSMC2 [11].

**Figure 3.**
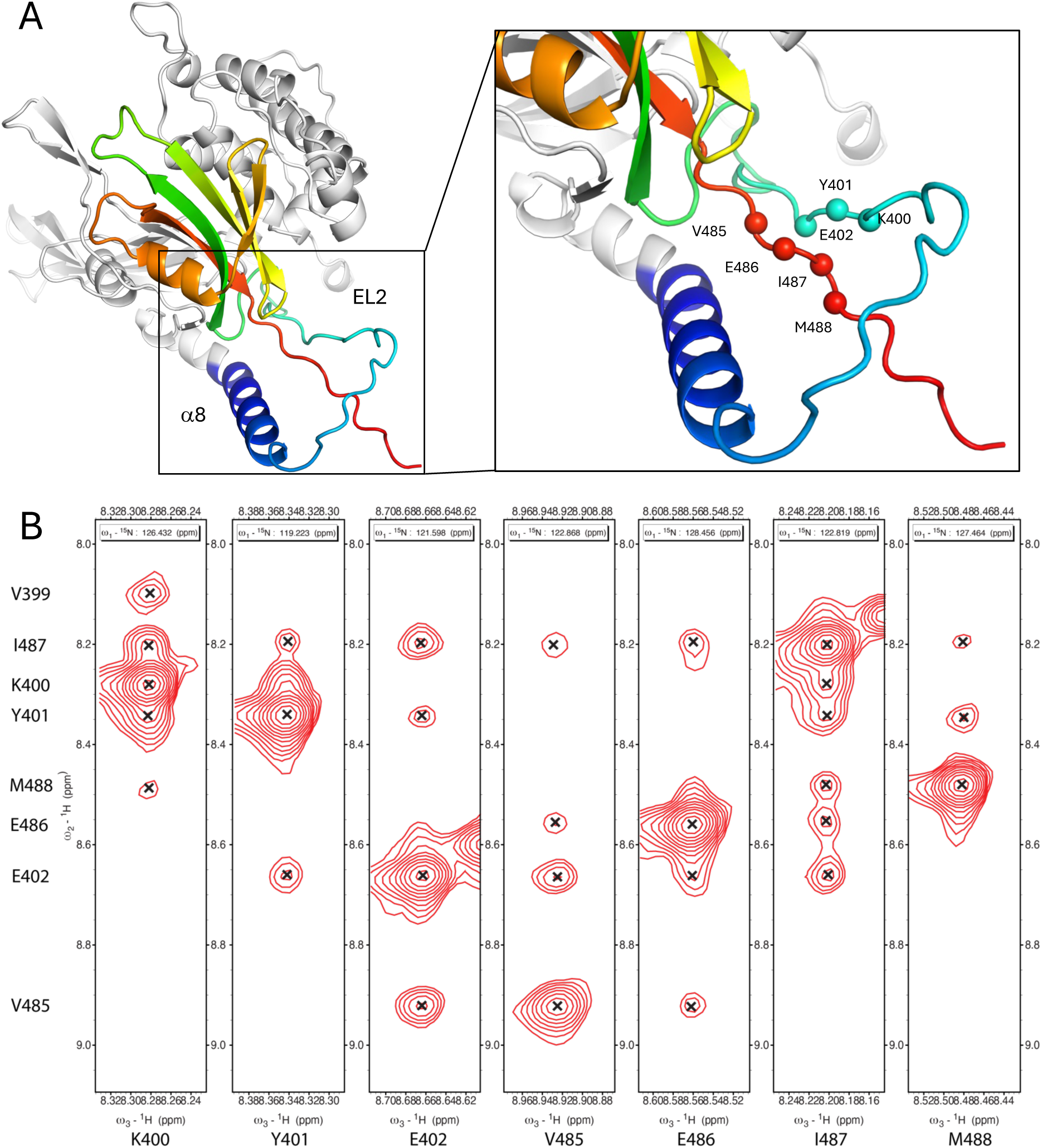
C-terminal tail in contact with EL2. (A) AlphaFold model of USP14_USP_ (with residue 368-494 in rainbow from blue to red) helix alpha 8 extending into EL2 wrapping the C-terminal tail. Highlighted with spheres for backbone amide nitrogen’s are residues 400-402 and 485-488. These residues show NOEs in the NOESY spectra in B. (B) Strip plots for ^15^N-edited NOESY TROSY-HSQC for residues 400-402 and 485-488.

By the CheSPI analysis, we were able to further detail the solution population of secondary structure in critical functional elements of USP14 including the four-stranded β-sheet (“fingers”) (βF1-2) and blocking loops BL1-BL3. In solution, βF1-2, where Ub rests in the USP14-Ub complex is well defined with high β-propensities for all four strands (β1-β2, β4-β5), whereas in the USP14-apo crystal structure, the outer strands of this sheet are less well resolved (β2 and β4) **(Figure 2A).** In solution, BL1 transiently forms a β-finger, with maximum β-strand populations of 20% (β7) and 60% (β8). Such β-finger propensity is less evident in USP14_USP_ crystal structures (PDBID: 2AYO, 2AYN) but more distinct in the yeast homologue Ubp6 (PDBID: 1VJV) and clearly present when USP14 is bound to the proteasome[11]. BL2 maintains partial β propensity up until residue pair G428/G433 and may thus have some increased β-structure propensity compared to crystal structures; lack of assignments at the tip precludes further analysis. Our data suggest that BL3 does not show regular secondary structure in solution, which agrees with the crystal structures and when USP14 is bound to proteasome. [11–13]

### NMR relaxation shows a continuous community of dynamic loops and reveals intersegmental dynamic connections

To extend our understanding of internal motions in USP14, we evaluated backbone ps-ns dynamics in USP14_USP_ (**Figure 2B, Supp Figure 2**). NMR relaxation properties were determined using ^1^H-^15^N TROSY-HSQC based longitudinal relaxation rate (R_1_) and transverse relaxation rate (R_2_) (800 and 900MHz) as well as steady state heteronuclear {^1^H}-^15^N NOE (hetNOE) experiments (900MHz) [42]. NMR dynamics evaluation is often interpreted by the Lipari-Szabo model-free formalism to quantify the global rotational correlation time (ρ_c_), the local order parameter (S^2^) and the R_ex_ contribution to relaxation due to slower motions, also called chemical exchange [43,44]. However, this simplified formalism often used for small proteins has been shown to be overgeneralized for large proteins at higher magnetic fields[45]. Since the molecular weight of USP14_USP_ (46 kDa) required higher magnetic fields for sensitivity and resolution, we therefore based our analysis on interpreting the relaxation experiments as such.

As expected, segments that follow the same dynamics as the core fold, within one standard deviation of the trimmed average across the three relaxation experiments, are predominantly located within secondary structure elements (**Figure 2)**. Higher R_1_’s, lower R_2_’s and/or hetNOE values below 0.65, typically found in loops, indicate residues with high local atomic fluctuation and increased backbone flexibility in the ps-ns range. Large motions and slower dynamics (µs-ms) generate complicated relaxation patterns and in many cases severe line broadening and thereby loss of peaks. Our data reveal such more complex patterns for USP14_USP_, both in loops and in secondary structured segments as determined by crystallography[12] and predicted by Alphafold[46] (**Figure 2).**

The most flexible regions in USP14_USP_ are EL1 and EL2 (**Figure 2, Supp Figure 2**). Dynamically, EL1 starts at Glu219 and re-joins the core fold dynamics at S241, entering a small helix that connects EL1 with βF1 in the finger region (**Figure 2**). Relaxation data supports dynamic closure of the EL1 loop by a stable Ile218 - Ile410 interaction, which is structurally observed in both crystal structures and in the AlphaFold model [38]. EL2 is part of the loop arrangement which wraps around the USP C-terminus (**Figure 2,3A**), which starts with a progressively flexible helix (α8), continues into the highly flexible EL2 and then anchors to the C-terminus at residues 400-402/485-488, as shown by distinctly reduced dynamics in both 400-402 and 485-488. After leaving the loop, the C-terminus is highly flexible **(Figure 2).**

When mapped onto the USP14_USP_ structure, the NMR relaxation data reveal a dynamic continuous community of loop segments with hetNOE values below 0.5, but not as extreme R_1_ and R_2_ values as EL1 and EL2 (**Figure 2D**). This community comprises the SL and structure elements anchoring onto the “catalytical” C114-containing helix, reaches out to the tip of the β-fingers, and includes the well-known BL1-BL3 and PKL loops. Within this community, the relaxation experiments suggest several different regimes of flexibility. The EL1 and EL2 dynamic loops are not part of this continuous surface, as they extend to the other side of the USP core fold. SL and PKL do not show a clear trend of higher or lower R_1_ or R_2_ values, suggesting significantly lower flexibility than the EL loops (**Figure 2B,2D**). Whereas BL1 shows rapid (ps-us) dynamics by higher R_1_s together with lower R_2_s and hetNOE values **(Figure 2B)**, the exchange broadening due to intermediate exchange in the tip of BL2 as mentioned earlier indicates slower dynamics (µs-ms) than in the BL1 loop, but still significantly more flexibility than in regions with defined secondary structure (**Figure 2B, 2D**). The R_1_ values are increased at the c-terminal part of BL3 and at the tips of βF1 and βF2, but no increase in R_2_ can be detected indicating increased backbone mobility but not as much flexibility as in the EL (**Figure 2B**).

In all three NMR experiments collected at 900MHz, we observe cases of divergent dynamics for single residues in otherwise stable segments, with relaxation values distinct from the core properties in at least two of the NMR relaxation experiments; these were also outliers in the 800MHz R_1_ and R_2_ data sets. Interestingly, such residues are often close to other flexible residues or segments (Y356/G46, T288/BL3) or to prolines, suggesting dynamic connections. The most extensive such cluster includes V120 (α1), A139 & L140 (surface loop), and A155, L156, R157, F160 and S162 (α3) (**Figure 2E**). Here, the corresponding side chains form an intersegmental connection extending from the surface and into the helix α1 that harbors the catalytic cysteine C114 and contacts the catalytic H435 and D451 (**Figure 2E**). Notably, α4, which is presumably in intermediate chemical exchange, could be part of this dynamic cluster, being sandwiched between the intersegmental connection and the flexible SL loop (**Figure 2E**).

### Single point mutations alter DUB activity with maintained structure and stability

Given that several of the identified dynamic regions in USP14 include residues that have been proposed as essential for USP14 function, we wanted to investigate whether the intrinsic dynamic properties of USP14 functionally contribute to catalysis and/or Ub binding. Proteasome binding is required to fully activate USP14 and has been suggested to involve a two-state loop model for BL1 and BL2 [12]. However, based on our NMR analysis and the dynamics present in these loops, the activation of USP14 cannot be explained by such a simple model. Assaying proteasome binding in solution by NMR is extremely challenging due to the high molecular weight of the system and has only been done for thermostable homologs to the catalytic 20S particle [47]. To resolve this, we set out to assay whether dynamic allosteric connections are present already in free USP14.

We first designed and functionally characterized a set of conservative or soft mutations with the aim to select mutants that by themselves affect USP14 function but not its structure, so as not to confuse altered dynamics with structural changes. We assessed how single mutations affected the DUB activity, overall structure and thermal stability of USP14_USP_ *in vitro*. In the BL2 loop, we chose the activating S432E mutant, to mimic phosphorylation [8]. For the active site, we selected the non-catalytic C114A mutant [9]. Additionally, we identified residues in the USP14_USP_ that interact with Ub (PDBID: 2AYO) and designed mutations aimed at reducing Ub binding at surface-exposed positions. These included D199A, E202K, Y333V, S431A, F331V, Y436F, D199A+S431A and F331V+Y333V. All mutants were cloned, expressed and purified as full length USP14 and in the same USP14_USP_ construct as used in the NMR relaxation analysis.

The activity of the full-length mutants was evaluated in two gel shift assays, one for binding and one for cleaving Ub (**Figure 4B,C,D**) [8,13] with wildtype (WT) USP14 as a positive control. The Ub-binding assay utilized the Ub-propargylamide probe. Upon Ub binding, the propargylamide moiety near the active cysteine form a covalent bond (**Figure 4B**). In the cleaving assay, USP14 was incubated with K48-linked di-Ub resulting in mono-Ub by active USP14 (**Figure 4C,D**). D199A and E202K eliminate activity in both the binding and cleaving assays, whereas Y333V reduced activity compared to the WT. To quantitatively compare deubiquitinating activity between WT USP14 and mutants D199A, E202K, Y33V, C114A and S432E, we conducted a Ub-rhodamine hydrolysis assay (**Figure 4E,F**). Consistent with the gel shift assays, the E202K and D199A mutants exhibited no hydrolysis activity, like the non-catalytic C114A mutant. On the other hand, Y333V and S432E demonstrated 20% and 430% of WT activity, respectively, providing additional evidence for the diminishing and augmenting effects of these single mutations on the DUB activity. Mutations on E202 and Y333 have previously been studied and shown to partially inactivate USP14 activity [11]. As a quality control for folding and stability, the USP14_USP_ mutants were analyzed with circular dichroism (CD) spectroscopy and differential scanning fluorometry (DSF) revealing highly similar secondary structure composition and stability (**Figure 4G, Suppl Figure 3**)

**Figure 4.**
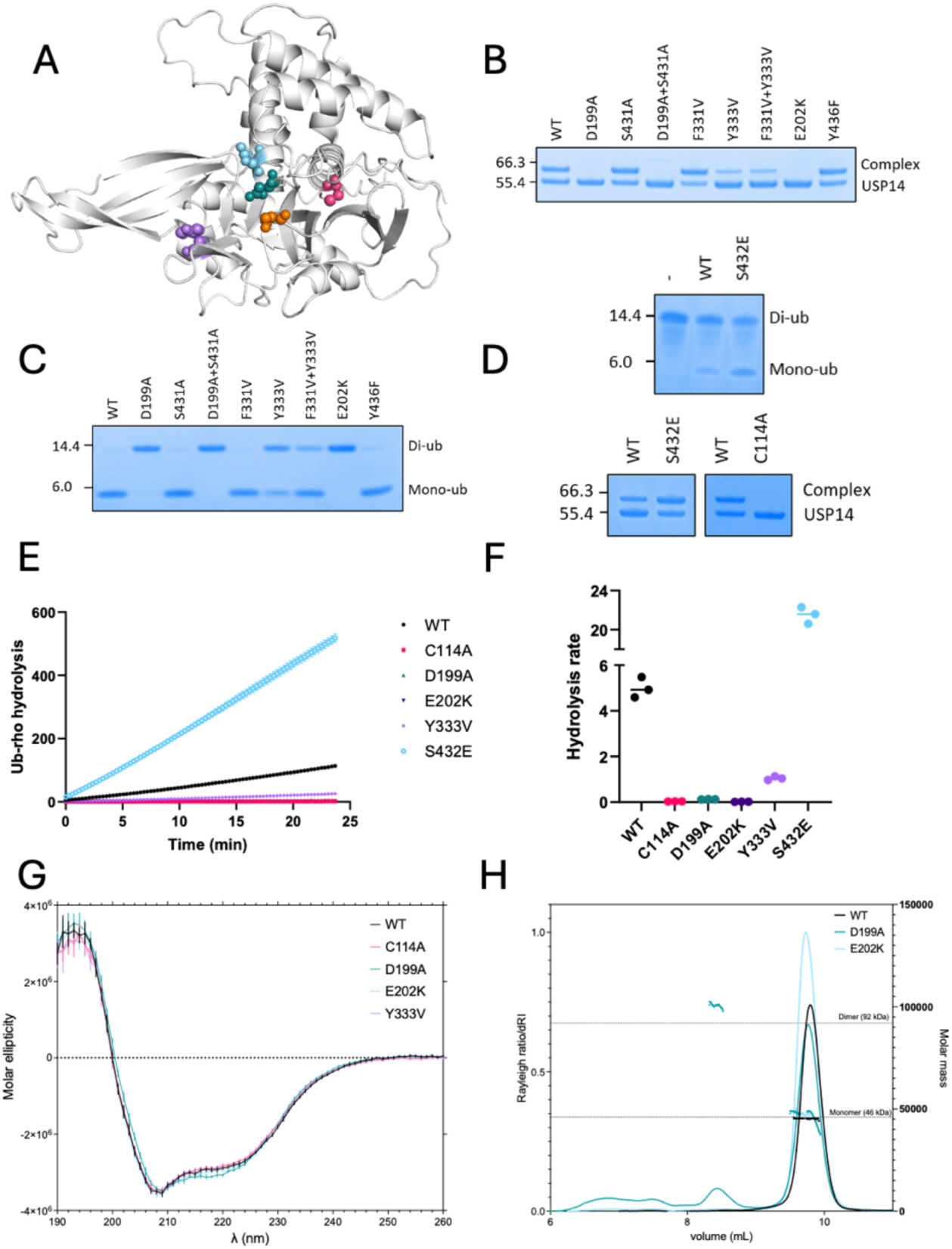
Evaluation of mutant activity. (A) Positioning of mutants shown on AlphaFold model C114A (pink), D199A (teal), E202K (light blue), Y333V (purple) and S432E (orange). (B) Ub-binding gel shift assay utilizing the Ub-propargylamide probe (8.5 kDa). When Ub binds into the binding pocket, propargylamide is positioned in proximity to the active cysteine and a covalent bond is formed, linking USP14 and Ub into a complex with higher molecular weight. (C) Ubiquitin-cleaving gel shift assay. USP14 was incubated with K48-linked di-Ub which is cleaved into mono-Ub by active USP14. (D) Ub-cleaving assay for S432E, and Ub-binding assay for S432E and C114A. (E) Representative results from Ub-rhodamine hydrolysis assay. (F) Ub-rhodamine hydrolysis reaction rates from three separate experiments. (G) Global secondary structure investigation using Circular dichroism on USP14_99-494_ wildtype (black), C114A (pink), D199A (teal), E202K (light blue) and Y333V (purple). (H) Oligomeric state detection using size exclusion chromatography multi angle light scattering (SEC-MALS) data on USP14 wildtype (black), D199A (teal), E202K (light blue)

To get first insights into the molecular effects of the mutations, ^1^H-^15^N TROSY-HSQC spectra were recorded for ^2^H-^15^N labeled WT USP14_USP_ and the mutants C114A, D199E, E202K, Y333V and S432E (**Figure 5**). Most of the resonances superimpose well with WT spectra for C114A, Y333V and S432E, which in agreement with results from CD and DSF suggests a retained fold with only minor structural and/or dynamic changes (**Figure 4G,H**,**Figure 5**). However, the D199A spectrum is severely disturbed with extensive loss of peaks, and the E202K spectrum lacks 20% of the resonances identified in the WT spectrum with 45% reduced peak intensity (**Figure 5, Suppl Figure 4**). Size exclusion chromatography coupled multiangle light scattering (SEC-MALS) measurements showed partial aggregation of D199A, with slight tendencies also for E202K at high MW (**Figure 4H**). Furthermore, while all mutants show two inflection points in the first derivative of the 350/330 nm ratio, the first inflection point for D199A significantly differs from the other mutants (**Suppl Figure 3A**). To focus on local structural and dynamic effects rather than larger-scale structural and possibly disruptive changes, we omitted the D199A and E202K mutants from further analysis.

**Figure 5.**
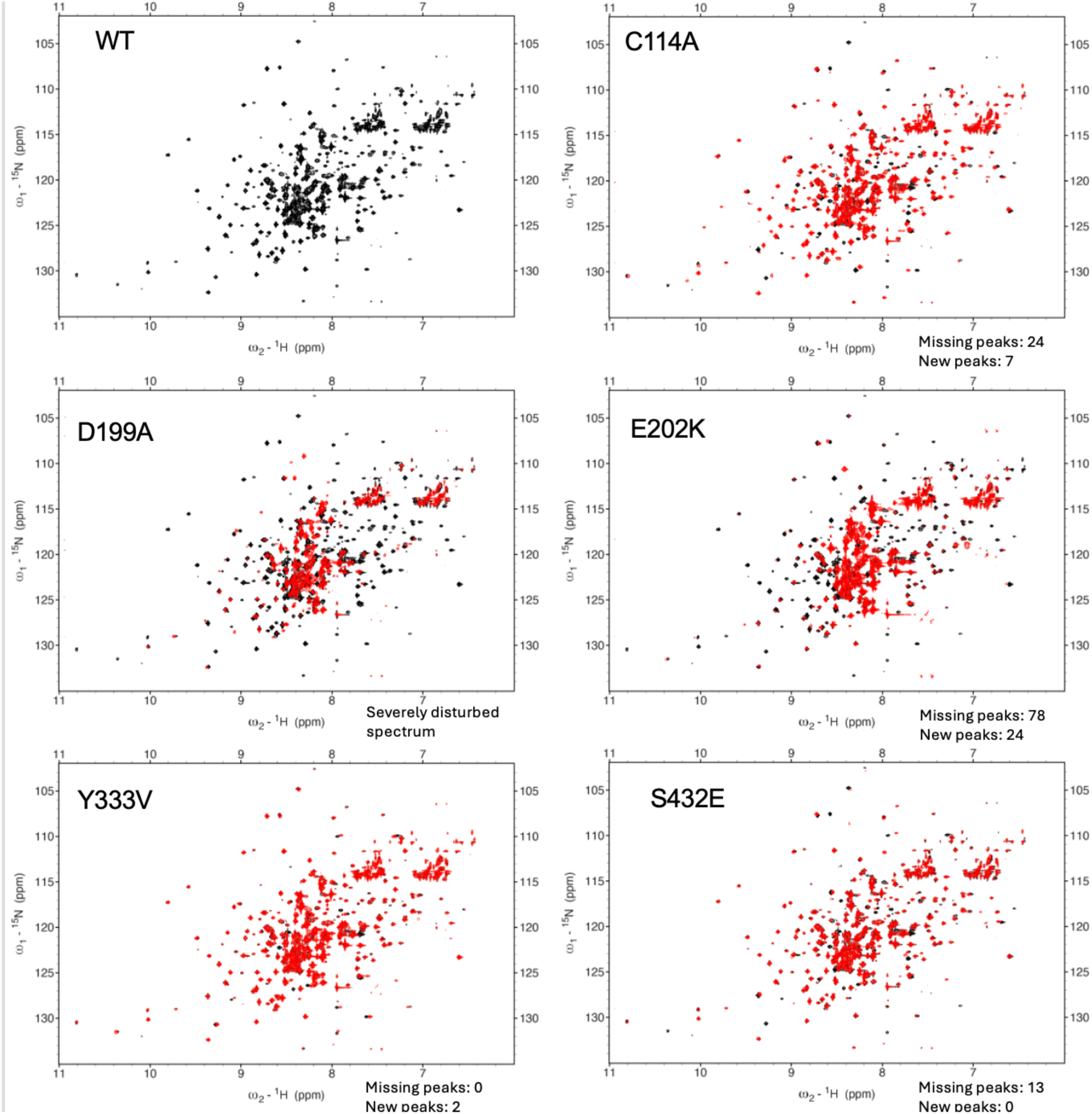
NMR characterization of USP14 mutants. ^1^H-^15^N TROSY-HSQC spectra USP14_99-494_ for mutants (red) overlayed on wildtype (black). WT spectrum on top left for reference.

### Interconnected dynamic networks in USP14 are affected by functional mutations

The NMR resonance chemical shift and line shape are both highly sensitive to small changes in structure and dynamics, thereby contributing a powerful tool to investigate dynamic networks in proteins [48,49]. The small spectral changes in the inactivating mutants (C114A, Y333V) and activating mutant S432E allowed us to readily assay CSPs and relative intensities between each mutant and WT USP14 **(Suppl Figure 4)** by direct comparison of TROSY-HSQC spectra (**Figure 5)**, using the NMR assignment of WT USP14.

For the catalytically inactive C114A mutation, significant CSPs are observed for 58 residues in several segments of USP14_USP_ **(Figure 5, Suppl Figure 4).** In addition to expected CSPs adjacent to the mutation site, CSPs of similar magnitude are observed at positions >10 residues apart from the mutation site in the SL, PKL, BL1, BL2 (including the second catalytic residue H435) and BL3 loops as well as and in the β-turn connecting strands β11-β12, housing the third catalytic triad residue D451 (**Figure 5, Suppl Fig 4).** In addition, peaks corresponding to 24 residues assigned in WT-USP14 were not identified in the C114A spectra, indicating dynamic line broadening or larger CSPs than would motivate deductive assignments [27]. Since only seven new peaks were detected, we assume most of the peaks that disappeared have been line broadened beyond detection, indicating a change in dynamics or/and multistate properties rather than in structural properties.

For the mainly inactivating Y333V mutant positioned in BL1, all residues could be assigned **(Figure 5).** Similar, but less extensive, network effects were observed as in the C114A mutant despite the surface-exposed structural position of this mutation, designed to interfere with Ub interaction. The highest CSPs were observed in the BL1 and BL2 loops, but also in the SL across from the groove binding the Ub c-terminal tail, and in the structurally more distant PKL and BL3, all part of the community of dynamic loops (**Figure 2D**). Distinct from C114A, no significant CSPs were observed for the active site residues C114, H435 and D451 **(Figure 5, Supp Figure 4)**.

Interestingly, the activity <u>enhancing</u> S432E mutant [8] affects similar loops and regions of USP14 as the inactive C114A mutant, including BL1, BL2, BL3, PKL, the loop preceding helix α1, and helix α1 comprising C114 but with less magnitude **(Figure 5, Supp Figure 4)**. The catalytic triad residue H435, which is well-ordered with little dynamics in the WT relaxation measurements, could not be assigned in this mutant, indicating a significant structural and/or dynamic perturbation. In contrast, no perturbation on C114 or D451 is detected in the activating mutant. Previous studies suggest that phosphorylation on S432 and the mutation S432E rearranges BL2 to a more open configuration, hence increasing USP14 activity [8]. Our results rather suggest more extensive dynamic and/or structural effects connected to the catalytic triad, in a pattern that is related to that observed in the inactivating mutation C114A.

The effects of the single mutations assayed by NMR suggest that larger, but specific, networks of residues are consistently affected by mutations affecting USP14 activity. To better understand the structural context of the effects observed by NMR in the functional mutants, we mapped residues with CSPs and/or line broadening effects onto the USP14 structure (**Supp Figure 4**). The perturbed regions in mutants C114A, S432E and Y333V are highly similar, with residues in the SL, BL1, BL2 and BL3 loops being perturbed by all three mutations (**Supp Figure 4**), forming an interconnected network that includes residues up to 25 Å away from the mutation site. This network of residues corresponds well with part of the community of dynamic loops identified by NMR relaxation (**Figure 2D**). However, regions of USP14 remain entirely unperturbed, including βF1-βF2, the domain core and loops located in other parts of the protein, including EL1 and EL2. Furthermore, although all three mutation sites are fully (S432E and Y333V) or partly (C114A) surface exposed, CSPs resulting from the mutations are consistently observed also for buried residues **(Supp Figure 5A)**. CONSURF analysis for USP14 based on PDB structures 2AYO and 2AYN show that this continuous, partly buried and partly surface-exposed region, is highly conserved, suggesting functional importance **(Supp Figure 5B)**.

### USP14 dynamic networks reach the proteasome interface, creating allosteric paths for reciprocal activation

USP14 enzymatic activity is enhanced by binding to the proteasome, and reciprocally, USP14 binding to the proteasome allosterically progresses proteasome activity [11,18]. Indeed, we found that USP14 regions affected by activating/deactivating mutations overlap significantly with regions partly or fully buried on binding the proteasome **(Figure 6**). Several residues within the BL1, BL2 and BL3 loops, which were consistently affected by the C114A, S432E and Y333V mutations but sufficiently far from the mutation site to not be directly affected, were also partly or fully buried upon proteasome binding. The PKL touches the proteasome-USP14 interface and was affected in the deactivating, but not in the activating, mutant. In contrast, the switching loop (SL) and the active site residues (C114, H435 and D451) were not part of the proteasome interface, and the perturbations resulting from the Y333V mutation did not reach the catalytic triad. Distinct from these effects, the EL2 loop is partly buried in several of the proteasome-bound USP14 conformations but is not affected by any of the mutations affecting USP14 catalytic activity as analyzed here **(Figure 6).**

**Figure 6.**
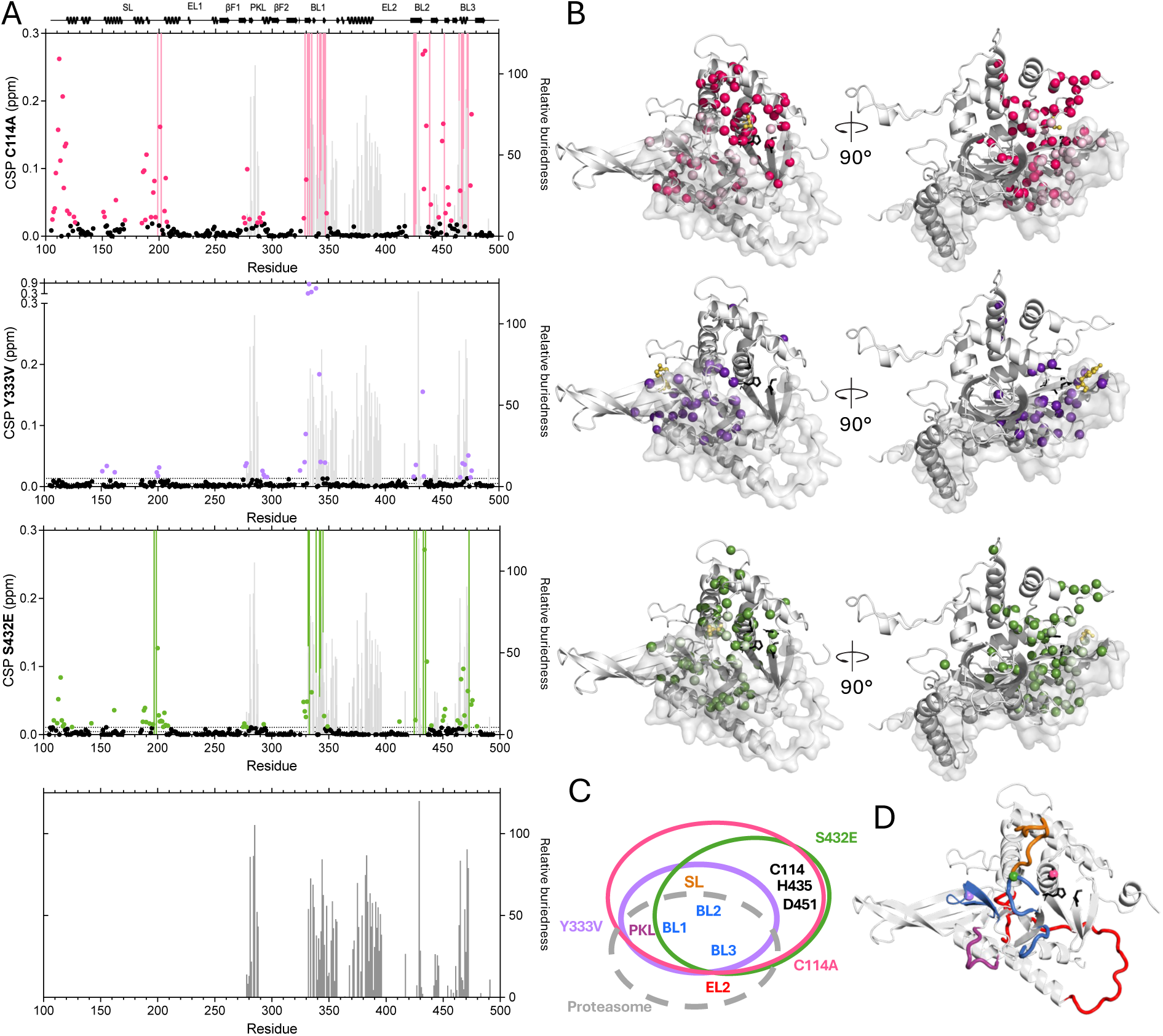
Joint and discrete USP14 perturbations and their allosteric connection to the proteasome interaction region. (A) CSPs are plotted as a function of sequence (dots), significant CSPs are highlighted with mutant specific colors: C114A (magenta), Y333V (purple), S432E (green). Missing residues are shown as mutant-specific colored bars. The relative buriedness (see methods) for all USP14 residues in all conformations identified in recent cryo-EM USP14-proteasome structures compared to free USP14 [11] are shown as gray bars. (B) Residues with significant CSPs are shown as magenta (C114A), purple (Y333V), or green (S432E) spheres. The proteasome interface is represented in gray surface. Mutation site shown as yellow spheres and catalytic triad as black sticks. (C) Venn diagram for the allosteric connections in USP14 detected by the CSPs of the different mutants C114A (magenta), Y333V (purple), S432E (green).

The regions affected by the functional mutations overlap well with the community of flexible loops and intersegmental dynamic connections identified by the NMR relaxation analysis (**Figure 2**). This suggests that the mutational effects are progressed through these dynamic networks in an allosteric manner, connecting the active site with the proteasome binding surface. Supporting the identification of allosteric networks, although the mutations were designed not to disturb structure or stability of USP14, CSP effects are not restricted to surface exposed loops but include buried residues in all affected areas of the USP14_USP_, as in other well-studied enzyme allosteric networks [21,23,49]. Assuming that these networks can signal the substrate engaged state of the USP14 active site to the proteasome, it is highly likely that the same networks can be engaged by proteasome binding at the other “physical end”, thereby activating USP14 catalytic and regulatory functions. Finally, the mutations do not reveal stabilisation of single states, rather the opposite as judged by line broadening effects, suggesting that the allosteric signalling path between the active site and the proteasome binding site is dynamic, including the BL1 and BL2 loops previously suggested to govern USP14 activity in an open/closed manner. Taken together, our work suggests that rather than stabilizing or destabilizing the conformational state of specific loops, the activation of USP14 by the proteasome is dynamically conveyed through distinct allosteric networks comprising interconnected flexible loops.

## Discussion

In this study, we conducted a comprehensive analysis of the catalytic USP domain of USP14 using NMR to map its dynamic and structural properties in solution. Our results consistently demonstrate the highly dynamic nature of loop communities in USP14_USP_ (BL1, BL2, BL3, SL, PKL) on the ps-ns timescale, as evidenced by NMR relaxation experiments (R_1_, R_2_, hetNOE). This implies that the USP14 loops can access a broad conformational space, encompassing both “open” and “closed” states. Extending this, we observe slower dynamics in the intermediary exchange regime as evidenced by line broadening and signal loss, particularly in the tip of the BL2 loop and helix α4. Furthermore, we show that in USP14, regions with slower dynamic properties consistently align with residues exhibiting faster dynamics, thereby linking motions in different regions and on different time scales. Our NMR mapping of the effects of USP14 functional mutations span over 45 Å and engage the community of loops and intersegmental connections forming dynamic networks that reach and include the proteasome-binding interface. This novel finding reveals the possibility that allosteric networks connect the USP14 active site with the proteasome interaction surface, and that these networks are pre-existing and activated by regulatory interactions.

Our direct experimental observation of extensive dynamics in the BL loops of free USP14_USP_ suggests that steric blocking of substrate binding by these loops is unlikely to be the main mechanism for USP14 autoinhibition. Instead, the activation of USP14 by proteasome binding may be allosterically conferred, resulting in altered dynamic properties of the substrate-engaging loops and within the active site region, rather than in distinct structural shifts as proposed from cryo-EM and crystallography[11,12]. This agrees well with the concept of triggering and triggered loops put forward for lipases and enolases, where coupled motion can contribute to long-range modulation of functional loops (reviewed in [20,50]). For USP14, the most efficient lead molecules in the IU1 series [9] predominantly target the opening of the active site groove, proposed to sterically prevent the access of the c-terminus of ubiquitin to the active site. However, these molecules show relatively weak activity in cell-based studies and do not inhibit USP14 non-catalytic proteasome regulating function (reviewed in [13]), suggesting the presence of more complex, allosteric modes of regulation.

The flexible properties we observe in the active surface loops, including BL1, BL2 and SL, support a dynamic functional model for USP14, which aligns well with an increasing body of research on USPs that supports the flexibility of BL loops. Recently, molecular dynamics (MD) simulations revealed differences in dynamics in BL1, BL2 and SL depending on the starting structure as well as on the phosphorylation state of S432 [51]. In USP15 crystal structures, both open and closed states of BL1, BL2 and SL loops have been observed in the absence of Ub, suggesting inherent dynamics [52]. In USP7, BL1, BL2, BL3 and SL regions were all identified as highly dynamic in MD simulations, both in the absence and presence of inhibitors binding into the Ub tail pocket [53]. Furthermore, although crystal structures captured free USP12 with a catalytic cleft obscured by collapsed BL1 and BL2 loops, biochemical data confirmed that free USP12 is fully capable of engaging its substrate, leading the authors to suggesting renaming these loops as binding loops rather than blocking loops [54].

Recent cryo-EM studies [11] show that the USP14 highly dynamic EL2 loop characterized in this work contributes most of the interaction surface to the proteasome AAA domain, which in turn alters position relative USP14 during the proteasome catalytic circle. However, EL2 is not affected by mutations in or close to the USP14 active site. The extensive EL2 flexibility, fine-tuned by the connection between EL2 and the C-terminus, may contribute enough plasticity for USP14 to remain in contact with the AAA domain throughout the different states of proteasome catalysis, while at the same time keep the USP-OB interaction through BL1-3 optimized for optimal loop position and dynamics for USP activity.

In this work, we identified by NMR relaxation a intersegmental connection reaching from residues in α3 into the buried, Cys114-carrying α1. Furthermore, this connection lies adjacent to helix α4 in USP14, which we cannot observe due to extensive line broadening. Although this USP14 region has not previously been assigned functional relevance, the helix corresponding to helix α4 in USP12 adopts shifted positions in different crystals structures, suggesting dynamics [54]. We note that this region corresponds to one of the dynamic USP touchpoints of the USP14_Ubl_ in full-length USP14 as determined through joint probing by NMR and SAXS techniques[39]. The possible functional relevance of this intersegmental connection will need to be further investigated.

The possibility that USP14 activity is jointly regulated by differently inducible, but partly overlapping allosteric networks installing communication between the active site and the spatially distinct Ub and proteasome binding sites is thus supported by observations in several other USPs. In USP12, two interacting proteins activate distinct yet complementary allosteric networks: UAF1 binding allosterically affects SL, while WDR20 binding allosterically activates the USP12 active site. [54]. In USP7, full activation of USP7 requires two intramolecular allosteric interactions: by its C-terminal tail, which dynamically binds the SL loop [55] – and by USP7 Ubl domains 4-5 which dynamically bind at similar USP sites as where the proteasome interacts with the USP14_USP_ [55,56]. In SAGA, full activation of Ubp8 (yeast analog to human USP22) requires two allosteric regulatory touch points: Sgf1 zinc finger contacts with the loop that houses the Ubp8 active-site cysteine, and SAGA modules Sgf1 and Sus1 occupying the same molecular space as the proteasome-interacting EL2 region in USP14 [57]. Indeed, multiple, interconnected allosteric networks could be an intrinsic property of this protein family.

Our findings reveal various aspects of dynamic connections within USP14, suggesting the presence of allosteric networks that link enzyme activity with regulatory functions. These allosteric networks, which include residues in the blocking loops, are interconnected through common residues, a phenomenon known as “multistate allostery”. The concept that USP14USP allosteric networks are pre-existing, coupled, and activated by regulatory interactions with the USP fold could be crucial for future targeted drug design.

## Methods

### Protein production

Perdeuterated 6xHis-tagged USP14_91-494_ and USP14_99-494_ were expressed in M9+ medium enriched with ^15^N-ammonium and deuterated ^13^C-glucose (USP14_91-494_ only) according to protocol developed by M. Cai et. al. [58]. The culture was induced by 0.5 mM IPTG and harvested after 20h at 25°C. The pellet was resuspended in 20 mM HEPES pH 7.5, 500 mM NaCl, 10 mM imidazole, 5% glycerol, 0.5 mM TCEP, 5 units/ml recombinant DNAse I and one EDTA-free protease inhibitor cocktail tablet per 75 ml and lysed by cell disruption at 25 psi in 4°C. Protein was purified as previously described [59] but with a Hiload 16/600 Superdex 200 pg column (GE Healthcare) for size exclusion chromatography (SEC). No unfolding and refolding of USP14 to optimize hydrogen exchange was pursued during purification, due to significantly reduced yields during refolding.

### NMR spectroscopy

Backbone assignment data were collected on Bruker Avance III HD 800 MHz spectrometer equipped with 3 mm TCI cryoprobe at the Swedish NMR centre. ^2^H, ^13^C, ^15^N labeled USP14_91-494_ were concentrated to 0.47 mM, in 20 mM HEPES, pH 7.5, 100 mM NaCl, 0.5 mM TCEP, 0.02 mM NaN_3_ and 10% D_2_O. All spectra were processed with NMRpipe [60] or mddNMR [61]. Backbone resonance assignment experiments were conducted at 30°C for USP14_91-494_ using TROSY based HSQC, HNCA, HNcoCA, HNCACB, HNcoCACB, HNCO, HNcaCO, ^15^N edited NOESY-TROSY-HSQC experiments and resonances were assigned by combining FLYA [62], COMPASS [63] and manual assignment in NMRFAM-SPARKY [64]. Complementary assignments for loops were achieved by combining the new experiments with spectral information from the ^13^C,^15^N labeled USP14_1-494_, where crowding was less of an issue. [39] CheSPI [40] was utilized for secondary structure populations based on N, H_N_, C_α_, C_β_, and C’ chemical shifts.

TROSY-HSQC were recorded for USP14_99-494_ WT and mutants according using 32 transients at 25°C with the concentrations 0.89 (WT), 0.53 (C114A), 0.28 (D199A), 0.89 (E202K), 1.07 (Y333V) and 0.58 (S432E) mM.

Peak intensity analysis of USP14_99-494_ mutants was performed by calculating the intensity ratio between the mutants and WT after normalization for concentration. CSPs between mutant and WT USP14_99-494_ were calculated according to

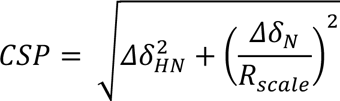

with the scaling factor R_scale_ = 6.5. The trimmed mean CSP and corresponding standard deviation (*σ*) as shown in Figure 3 were calculated stepwise by first calculating the mean and *σ* of the whole data set, after which any value exceeding one *σ* was excluded and a new mean and *σ* was calculated for the remaining values.

### Relaxation data collection and analysis

TROSY-optimized ^15^N R_1_, R_2_ and hetNOE experiments [42] were collected as pseudo-3D at 25°C using a Bruker Avance III HD 900 MHz spectrometer equipped with 3 mm TCI cryoprobe at the Swedish NMR centre, on USP14_99-494_ with a concentration of 0.89 mM in same buffer as above. For the relaxation measurements, the USP construct was slightly shortened in the N-terminus, compared to the assignment experiments, to exclude the flexible region 91-98 [39]. The R_1_ and experiment were collected using 12 different relaxation delays and 2 duplicates: 0.08, 0.08, 0.16, 0.24, 0.32, 0.4, 0.4, 0.56, 0.72, 0.8, 1.04, 1.2, 1.6 and 2.4 s and the R_2_ experiment with 10 different relaxation rates, one duplicate and one triplicate: 0.008, 0.016, 0.016, 0.016, 0.031, 0.047, 0.063, 0.094, 0.110, 0.110, 0.125, 0.156, 0.203 and 0.250 s for R_2_. The steady-state hetNOE values were determined from the ratios of the average intensities of the peaks with and without 5 s proton saturation.

The NMR relaxation data was analyzed using the software PINT [65] in which peaks were integrated, fitted with a combination of Gaussian and Lorentzian line shapes and relaxation parameters determined. Errors were determined from the jackknife approach for the R_1_ and R_2_ experiments while hetNOE errors were determined based on measured background noise levels using the following relationship [66]:

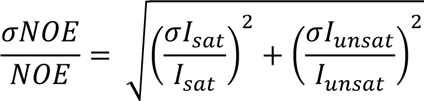

Residues with relaxation errors larger than 20% than the relaxation parameter were excluded from the dataset, together with residues with bad 3D fits or overlapping peaks. The trimmed mean and corresponding σ was calculated as for the CSP.

### Solvent accessibility calculations

Naccess [67] was utilized to determine solvent accessibilities for residues in the AlphaFold prediction model of apo USP14 and USP14 in complex with the proteasome (PDB-ID 7W3A, 7W3B, 7W3C, 7W3F, 7W3G, 7W3H, 7W3I, 7W3J, 7W3K, 7W3M), using the default probe size of 1.40 Å. For USP14 in complex with proteasome, the USP14 and Ub chains was copied to new PDB-files from each cryo-EM structure to only determine the binding interface between USP14 and the proteasome. Naccess was executed for both USP14+Ub and the full USP14+Ub+proteasome complex. The relative buriedness was obtained by subtracting the summed all-atom relative solvent accessibility of USP14+Ub+proteasome residues from USP14+Ub. Amino acids with relative buriedness > 10 was considered buried.

### Gel shift assays

Two gel shift assays were performed to screen the activity of the binding pocket mutants. For the screening, USP14_1-494_ was used. To test binding to Ub-propargylamide (Ub-PA), 3 μM USP14 was incubated with 6 μM Ub-PA (Biotechne, U-214-050) at room temperature for 24 hours in 20 mM HEPES (pH 7.5), 150 mM NaCl, 0.5 mM TCEP and 5 % DMSO. The same assay was also performed with Ub-vinylsulfone (Biotechne, U202-050) with the same results (data not shown), but the PA assay was chosen due to its lower reactivity which will require proper docking of ubiquitin in the binding pocket before covalent attachment. Di-Ub cleavage was tested by incubating 6 μM USP14 with 11.6 μM K48-linked di-Ub (Biotechne, UC-200B-025) for 24 hours at 30 °C in 16 mM HEPES (pH 7.5), 120 mM NaCl and 0.40 mM TCEP. Results were analyzed by SDS-PAGE with Coomassie staining. Three separate experiments were performed, and one representative gel is shown.

### DUB activity assay

Samples of 5 µM USP14 were prepared in triplicates in a black 384-well plate (corning 3820) with reaction volume 20 µl. The assay buffer contained 50 mM HEPES pH 7.5, 50 mM NaCl, 5 mM MgCl_2_, 1 mg/ml BSA, 1 mM DTT and 2 mM ATP. 1 µM Ub-rhodamine (Bio-Techne, U-555-050) was added to start the reaction. The hydrolysis of Ub-rhodamine was monitored with a Promega plate reader at 475 nm at 37 °C. The average background was subtracted. Three separate experiments were performed, the reaction rates in each experiment were calculated by simple linear regression and significance was evaluated by t-test in Graphpad Prism 10.

### Circular dichroism

Circular dichroism measurements were performed on a Chirascan spectrometer (Applied Photophysics) using a quartz cuvette with 1 mm pathlength. All samples were dialyzed against 10 mM phosphate buffer, pH 7.5 and centrifuged to remove aggregates. The wavelength scans were measured in samples containing approximately 3 μM USP14, determined by Nanodrop and theoretical extinction coefficients for each construct. CD spectra for USP14_99-494_ were recorded at 20 °C in the wavelength range 190-260 nm with 1 nm steps, 1.5 seconds per point and 10 repeats. Spectra for USP14_1-494_ were recorded with 2 nm steps. The average background of the dialysis buffer was subtracted, and the units were converted to molar ellipticity based on protein concentration measured in the sample by the DC protein assay kit (Bio-Rad) using WT USP14 as standard. Spectra for S432E was collected separately and the units were converted based on concentrations measured by Nanodrop.

### NanoDSF

NanoDSF was measured in the dialyzed samples prepared for CD spectroscopy. The samples were centrifuged to remove aggregates before the concentration was measured by Nanodrop (Implen) and they were then diluted to 1 mg/ml with 10 mM phosphate buffer pH 7.5. Unfolding was monitored in triplicates by Prometheus NT.48, in nanoDSF grade high sensitivity capillaries, with a temperature increase of 1 °C/minute. T_m_ values were calculated by the accompanying software and standard deviations are used as an error estimate.

### SEC-MALS

Multi-angle light scattering (MALS) analysis was performed following size-exclusion chromatography (SEC) using an in-line multi-detector instrument from Wyatt Technologies: consisting of DynaPro Nanostar and Mini-Dawn TREOS detectors coupled to an OptiLab T-Rex refractometer. 20 μl of USP14_99-494_ of either WT, E202K, D199A was injected onto a Superdex 75 Increase 10/300 analytical column (GE Healthcare) equilibrated in 20 mM HEPES (pH 7.5), 150 mM NaCl and 0.5 mM TCEP at a flow rate of 0.5 ml/min. The load concentration where 320 μM for WT, 470 μM for E202K, and 423 μM for D199A. Scattering data were used in combination with concentration estimates obtained from differential refractive index measurements (using a dn/dc = 0.185 mL/g) to deduce analyte molecular weight (MW). The measurements were done at 20°C. The MW distribution of species eluting from the column was determined using ASTRA7 software (Wyatt Technology).

## Acknowledgement

This research has received funding from the Swedish Research Council (M.S., P.D.), the Swedish Cancer Foundation (M.S., P.D.), The Swedish Childhood Cancer Foundation (M. S.), and the LiU Cancer research network (M.S., P.D.). We would like to acknowledge the ProLinC core facility, funded by Linköping University, for access to GE ÄKTA systems, DSF and CD spectrometer. We thank the Swedish NMR center for access to 800 and 900 MHz Bruker Avance III HD NMR spectrometers and Dr. Cecilia Persson and Dr. Ulrika Brath for their expert assistance in data collection. We acknowledge Professor Stig Linder, Professor Björn Wallner and Dr Vivian Morad for critical discussions concerning the research project.

## Supplementary Figures

**Supplementary Figure 1.**
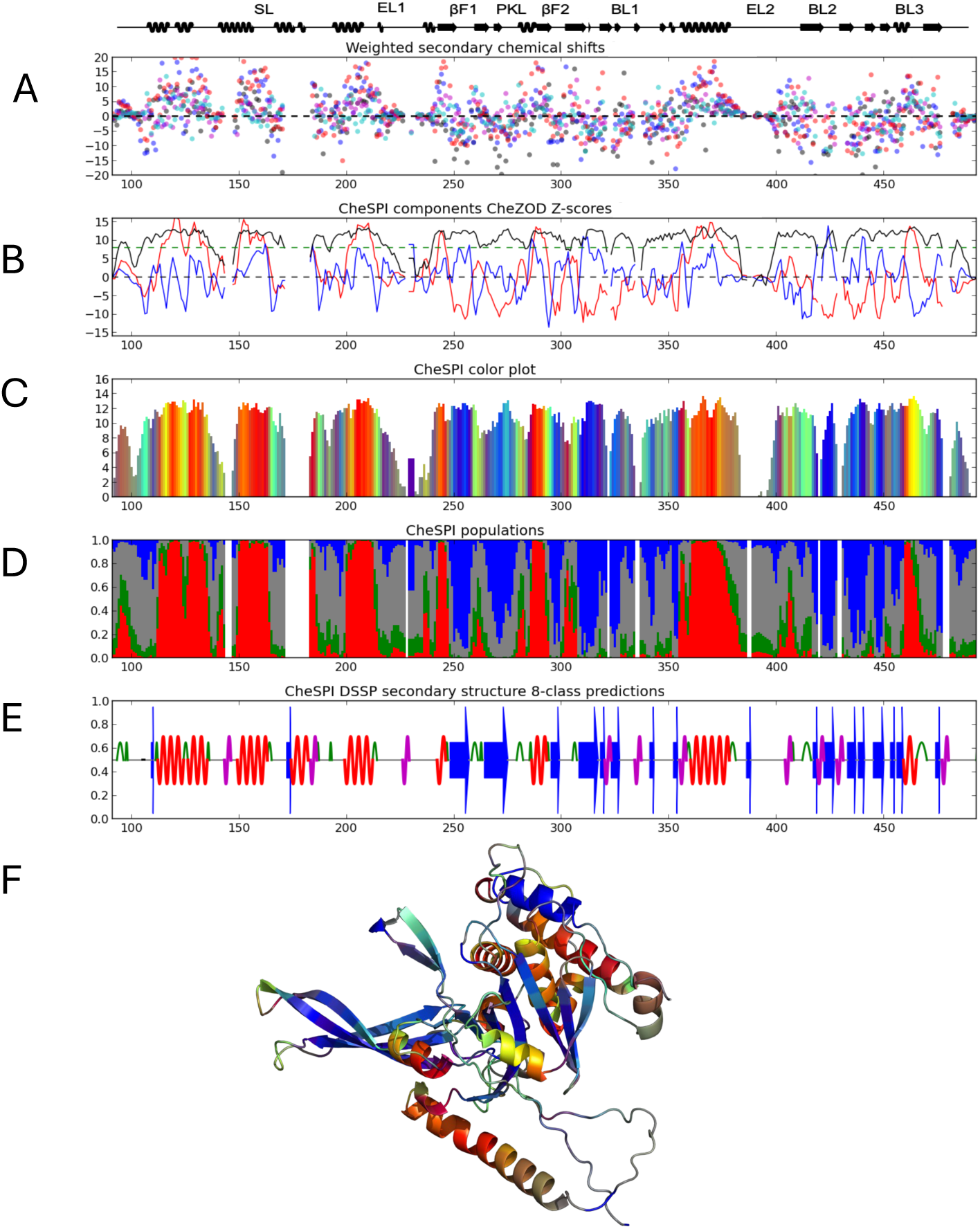
CheSPI results. Summary of CheSPI evaluations based on chemical shift data from USP14_USP_ (91-494), For reference the secondary structure elements from the Alphafold model is shown and annotated on top (A) Weighted difference between observed and predicted shifts shown with blue, red, black, cyan, and magenta dots for C’, C_α_, C_β_, H_N_, and N, respectively. (B) CheSPI components (Blue and red) and CheZOD Z-scores (black). Green dashed lines at Z = 8.0 for reference, CheZOD Z-scores < 8 are classified as disordered. (C) Bar plot colored according to the CheSPI color scheme. CheZOD Z-scores are used for bar heights. (D) Secondary structure populations as shown in Figure 2. (E) Illustration of the most confident secondary structure prediction. (F) A visual interpretation of the CheSPI result mapped onto the AlphaFold2 structure, extending the color scheme in A to provide an intuitive overview of the local structure and dynamics of UPS14 based on its chemical shifts **(Suppl Fig 1)**[40].

**Supplementary figure 2.**
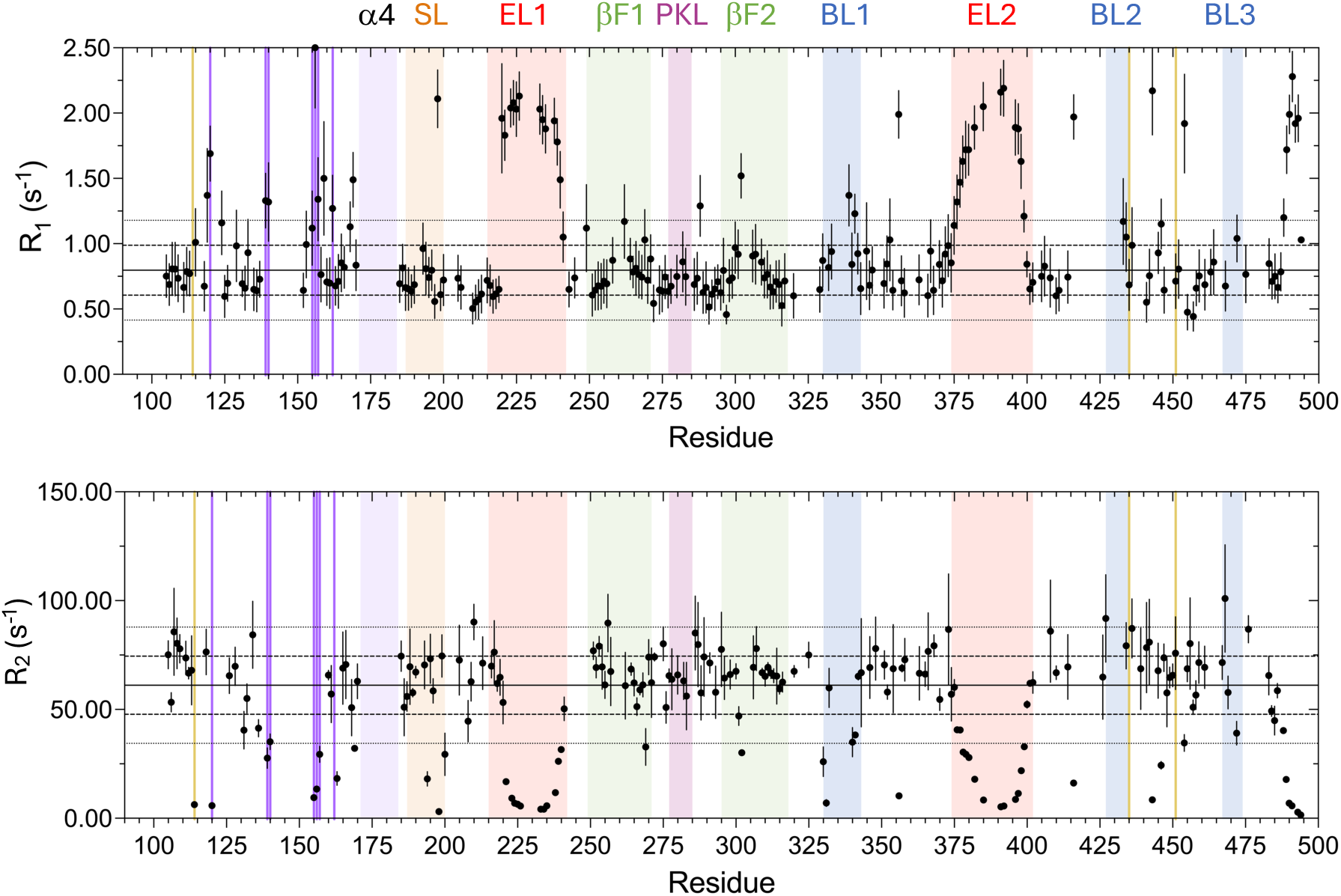
NMR relaxation data 800MHz. NMR relaxation data (R_1_ and R_2_ at 800MHz) for USP14_99-494_, revealing dynamic properties in the ps-ns range, as a function of sequence. For R_1_, R_2_ and hetNOE, the trimmed average value is indicated with a straight line. The values for one (dashed) or two (dotted) standard deviations above or below the trimmed average are indicated as lines. Color coding: Helix α4 light lilac, Switching loop (SL residues 188-199) in orange, Proximal Knuckle Loop (PKL residues 278-285) in purple, β-fingers in green (βF1 residues 249-272 and βF2 residues 295-319), and blocking loops (BL1-3) in blue (BL1 residues 330-342, BL2 residues 428-434 BL3 residues 468-473), Extended loops (EL1-2) (EL1 residue 214-242 and EL2 384-416) in red, dynamic cluster in dark lilac, catalytic triad C114, H435, D451 in yellow.

**Supplementary figure 3.**
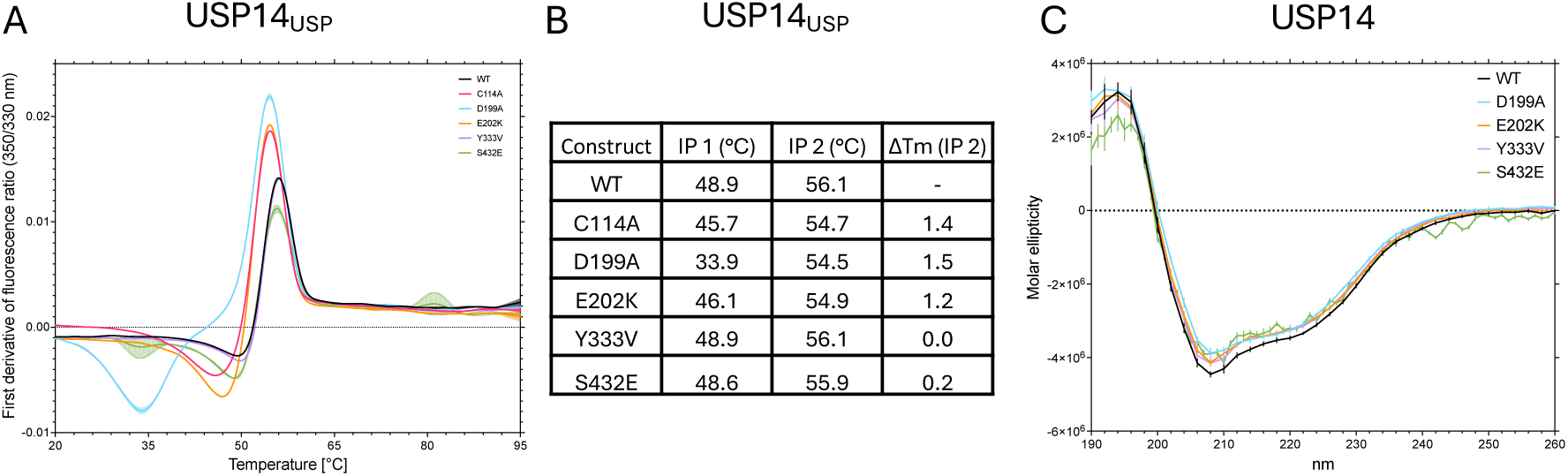
Supplementary evaluation of mutant activity. (A) First derivative of thermal stability of aromatic environment detected by differential scanning fluorescence (DSF) on USP14_99-494_ wildtype (black), C114A (pink), D199A (teal), E202K (light blue) and Y333V (purple). (B) Summary of DSF results. (C) Global secondary structure investigation using Circular dichroism on USP14_1-494_ wildtype (black), C114A (pink), D199A (teal), E202K (light blue) and Y333V (purple).

**Supplementary Figure 4.**
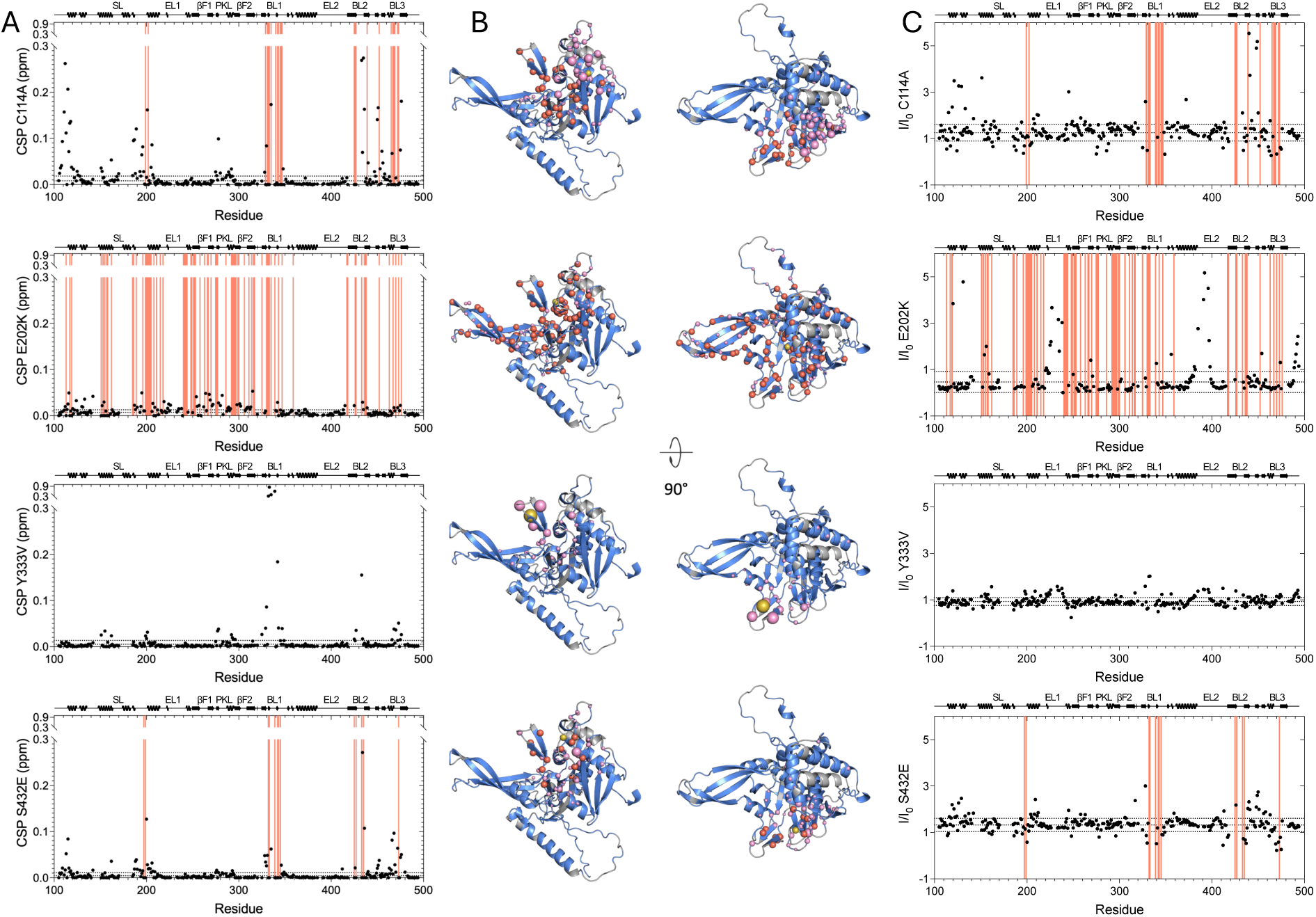
Chemical shift and intensity perturbations of USP14 mutants. (A) Chemical shift perturbations comparing USP14 WT and mutants (C114A, E202K, Y333V and S432E) plotted as a function of sequence. Red lines show missing peaks or peaks that have moved so much that assignment was not possible indicating large perturbations. (B) CSPs (pink) and missing peaks (red) from (A) is plotted on AlphaFold model. (C) Intensity ratios normalized by concentration between mutant (C114A, E202K, Y333V and S432E) and WT USP14. Red (See A)

**Supplementary Figure 5.**
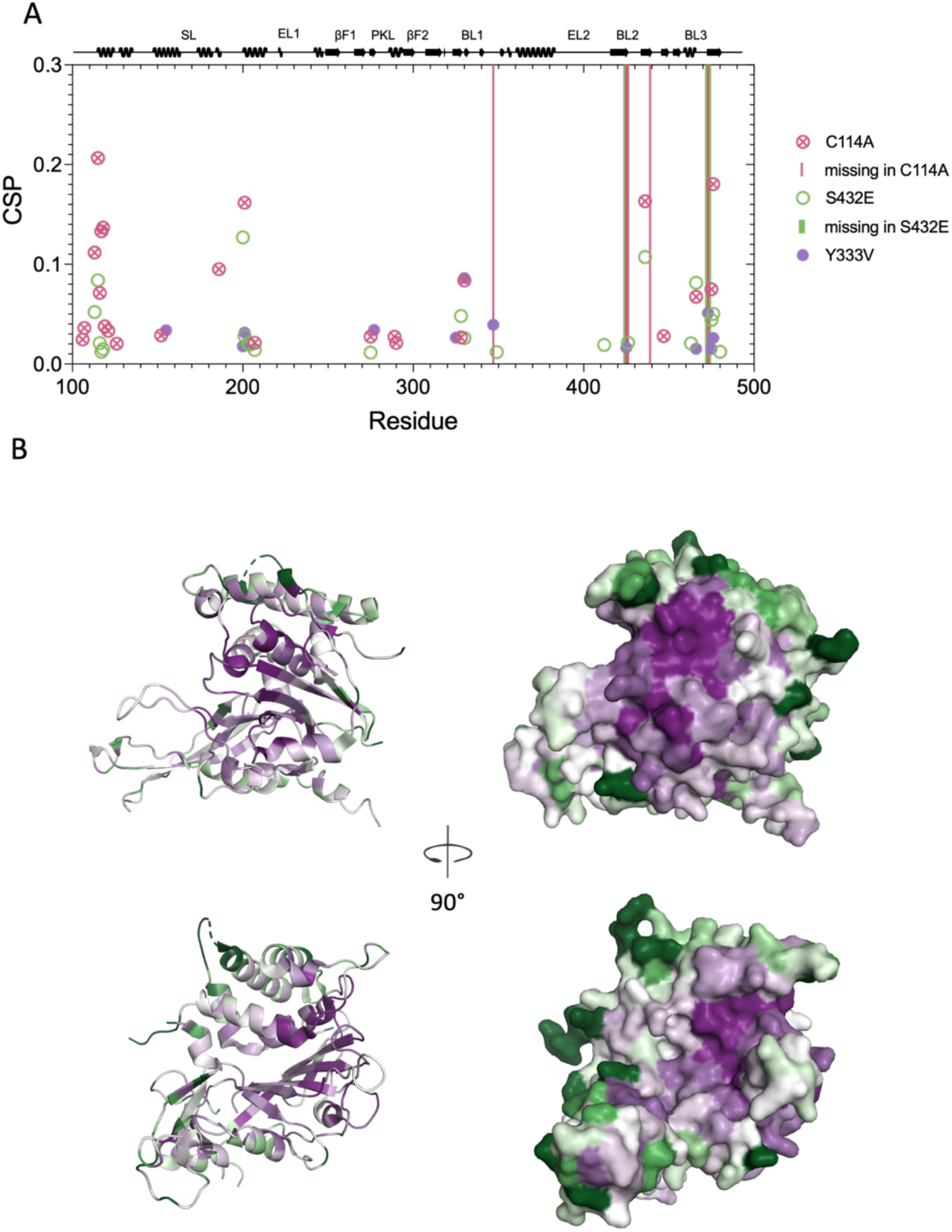
Allosteric networks coincide with conservation. (A) CSPs of residues in C114A, S432E and Y333V that have a total amino acid surface accessibility < 10 determined by Naccess in the USP14 AlphaFold model. (B) Consurf DB (consurf.tau.ac.il) visualisations of residue conservation mapped onto the USP14_USP_ crystal structure (PDBID: 2AYN, same result for 2AYO). The color scale goes from deep purple (highly conserved) over white to dark green (not conserved).

